# Antioxidant Gene Expression in Vocal Hindbrain of a Teleost Fish

**DOI:** 10.1101/354977

**Authors:** Clara Liao, Ni Y. Feng, Andrew H. Bass

## Abstract

Plainfin midshipman fish (*Porichthys notatus*) have a remarkable capacity to generate long duration advertisement calls known as hums, each of which may last for close to two hours and be repeated throughout a night of courtship activity during the breeding season. The midshipman’s striking sound production capabilities provide a unique opportunity to investigate the mechanisms that motor neurons require for withstanding high-endurance activity. The temporal properties of midshipman vocal behaviors are largely controlled by a hindbrain central pattern generator that includes vocal motor neurons (VMN) that directly determine the activity pattern of target sonic muscles and, in turn, a sound’s pulse repetition rate, duration and pattern of amplitude modulation. Of the two adult midshipman male reproductive phenotypes -- types I and II-- only type I males acoustically court females with hums from nests that they build and guard, while type II males do not produce courtship hums but instead sneak or satellite spawn to steal fertilizations from type I males. A prior study using next generation RNA sequencing showed increased expression of a number of cellular respiration and antioxidant genes in the VMN of type I males during the breeding season, suggesting they help to combat potentially high levels of oxidative stress linked to this extreme behavior. This led to the question of whether the expression of these genes in the VMN would vary between actively humming versus non-humming states as well as between male morphs. Here, we tested the hypothesis that to combat oxidative stress, the VMN of reproductively active type I males would exhibit higher mRNA transcript levels for two superoxide dismutases (*sod1*, *sod2*) compared to the VMN of type II males and females that do not hum and in general both of which have a more limited vocal repertoire than type I males. The results showed no significant difference in *sod1* transcript expression across reproductive morphs in the VMN and the surrounding hindbrain, and no difference of *sod2* across the two male morphs and females in the SH. However, we observed a surprising, significantly lower expression of *sod2* transcripts in the VMN of type I males as compared to type II males. We also found no significant difference in *sod1* and *sod2* expression between actively humming and non-humming type I males in both the VMN and surrounding hindbrain. These findings overall lead us to conclude that increased transcription of *sod1* and *sod2* is not necessary for combatting oxidative stress from the demands of the midshipman high-endurance vocalizations, but warrant future studies to assess protein levels, enzyme activity levels, as well as the expression of other antioxidant genes. These results also eliminate one of the proposed mechanisms that male midshipman use to combat potentially high levels of oxidative stress incurred by motor neurons driving long duration vocalization and provide more insight into how motor neurons are adapted to the performance of extreme behaviors.

## INTRODUCTION

Metabolic processes and cellular respiration produce highly reactive free radicals, capable of damaging many targets such as DNA, proteins, and nucleic acids, leading to cell damage. An accumulation of reactive oxygen species can be toxic when antioxidant defenses are insufficient to combat the rate of free radical production (Lobo et al., 2010). Oxidative stress has been highly implicated in a number of motor neuron diseases, including multiple scelerosis, Huntington’s disease, and amytrophic lateral scelerosis (ALS) (Adamcyzyk et al., 2016; Kumar et al., 2016; Bruijn et al., 2004). Neuroprotective strategies to combat oxidative stress-induced neurotoxicity have been reported, including the increased expression of endogenous antioxidants in regions of oxidative stress; for example, increased glutathione peroxidase is observed in the microglia of Parkinson’s Disease tissue (Peres et al., 2016; Pocernich 2000, Power et al., 2008).

Two enzymes, Cu/Zn superoxide dismutase (SOD1) and mitochondrial Mn-superoxide dismutase (SOD2), catalyze the reduction of the free radical O_2^−^_ and function as endogenous antioxidants to decrease neuronal oxidative stress. SOD1 is a crucial combatant to augmented oxidative stress that ensures neuronal survival, as SOD1 gene mutations have been implicated in the cause of motor neuron death observed in some familial cases of ALS (Bruijn et al., 2004). Higher expression patterns of aortic *sod2* during prenatal glucocorticoid exposure in monkeys also suggest that the mechanism of *sod2* upregulation serves as an important defense against oxidative stress (Atanasova et al., 2009)

As a model for investigating the potential role of antioxidants in motor neuron function, we studied the vocal motor system of the plainfin midshipman (*Poricthys notatus)*, a species of highly sonic teleost fish. Territorial males, also known as type I males, build nests beneath rocky shelters in the intertidal zone along the northwest coast of the United States and Canada during the late spring and summer and broadcast advertisement calls known as hums to attract females to their nest for spawning (Ibara et al., 1983; Brantley and Bass, 1994). Individual hums may last for up to two hours and are produced repetitively throughout an evening of courtship activity (Brantley and Bass, 1994; McIver et al., 2014; Feng and Bass, 2016). Type I males also produce repetitive series (trains) of two types of agonistic signals during nest and egg defense; brief (~100 ms on average) grunts occur at rates of 1-2 Hz and longer (~3 s on average) growls, a hybrid of hums and grunts, are generated at more irregular intervals (Brantley and Bass, 1994; McIver et al., 2014). An alternative, type II male morph neither builds nests nor courts females with hums; rather, they sneak or satellite spawn in an attempt to steal fertilizations from type I males. Type II males and females have only been observed to make irregular grunts (Brantley and Bass, 1994). The three reproductive morphs differ in a suite of somatic, neuroendocrine, and vocal motor traits, with a general trend that type II males are more similar to females (reviews: Bass, 1996; Feng and Bass, 2017).

A hindbrain central pattern generator (CPG) that exhibits extreme synchronicity and temporal precision in motor neuron firing (Bass and Baker, 1990; Chagnaud et al., 2011, 2012) drives the simultaneous contraction of a single pair of striated muscles attached to the walls of the gas-filled swim bladder (Cohen and Winn, 1967). The vocal CPG includes three topographically separate pairs of neuronal populations: vocal prepacemaker (VPP), pacemaker (VPN), and motor (VMN) neurons whose activity pattern directly translates into sound duration (VPP), pulse repetition rate (and fundamental frequency for multi-harmonic hums; VPN), and amplitude (VMN) (Bass and Baker, 1990; Chagnaud et al., 2011, 2012). VMN output (the paired VMN fire in phase) generates a vocal motor volley comprised of spike-like potentials that reflect the synchronous activity of VMN populations and are matched 1: 1 with each contraction cycle of the sonic muscles, resulting in one sound pulse (Cohen and Winn, 1967; Bass and Baker, 1990).

A recent study using next generation RNA sequencing (RNAseq) to investigate seasonal and daily variation in gene expression in the VMN of type I males revealed candidate genes proposed to underlie and support vocal CPG function (Feng et al., 2015). Many genes involved in cellular respiration pathways and several that code for antioxidant enzymes, including *sod1*, were expressed at higher levels in the VMN compared to surrounding hindbrain tissue. We proposed that these results were indicative of VMN’s high metabolic requirements and susceptibility to oxidative stress due to prolonged and high rates of cellular activity to generate vocalizations during the breeding season, in particular hums. These findings motivated the present study to investigate both *sod1* and *sod2* gene expression in the VMN and to compare it to the surrounding hindbrain across all three reproductive morphs – type I males, type II males, females – as well as between actively humming and non-humming type I males. We hypothesized that in order to combat oxidative stress, the VMN may (1) depend on higher *sod* expression in type I males that alone are known to generate long duration calls – hums, grunt trains, growl trains – as compared to type II males and females, and 2) depend on higher *sod* expression in type I males that are actively humming compared to those that are quiescent.

## MATERIALS AND METHODS

### Animals

For the sex/reproductive morph study, fish were hand collected from nest sites in northern California in June through August. This included fish from each of the three reproductive morphs: type I males (n = 10; median length: 14.95cm), type II males (n = 10, 9.15cm), and females (n = 8, 14.6cm). In the behavioral state study, type I males (n = 18, 23.55cm) were hand collected from nests in May through June from sites in both northern California and Washington state (see Bass, 1996 and McIver et al., 2014 for photographs of the sites). Fish were group housed near each of the collection sites for 2-3 days before shipping overnight to Cornell University, where they were housed individually in 10 gal (2015) or group housed in separate compartments of 100 gal (2016), artificial seawater aquaria at 16°C (close to natural habitat) prior to tissue collection (see Feng et al., 2016 for housing details). Fish were held on a 15h:9 h light: dark cycle to mimic the summer mating season photoregime (Feng and Bass, 2016). All fish were sacrificed within a month of collection.

### Behavioral State Study

For the behavioral state study, type I males were allowed to set up nests following the methods described in Feng and Bass (2016). Hydrophones placed near the nest were used to monitor vocalization activity. Males were allowed to hum for a minimum of 30 min prior to being anesthetized and sacrificed (see next section). Fish were anesthetized close in time as matched pairs of actively humming and non-humming individuals. To our knowledge, the non-humming fish were not making any other types of sounds (see Introduction). The non-calling fish were anesthetized and sacrificed within the same hour as the actively-humming fish to control for potential variation due to circadian rhythm. Brain tissue was immediately extracted in all cases.

### Tissue collection and processing

All fish were anesthetized in artificial seawater with 0.025% benzocaine prior to being killed by exsanguination from the heart. In each fish, the brain was surgically extracted, separated into forebrain-midbrain and hindbrain components by a transverse cut at the level of the cerebellum, and stored in RNAlater at −80°C. Immediately before RNA extraction, the VMN was surgically excised *in toto* from the surrounding hindbrain tissue using fine forceps (Fergus and Bass, 2013; Feng et al., 2015). RNA was extracted from VMN tissue and the surrounding hindbrain using glass homogenizers and TRIzol (Life Technologies) following manufacturer’s instructions. RNA was DNase-treated with Ambion DNase I (Invitrogen) and RNA concentration was assayed using Qubit fluorometric quantitation with the manufacturer’s RNA high sensitivity protocol (Life Technologies). cDNA was synthesized from DNase-treated RNA with Invitrogen Superscript III Reverse Transcriptase.

### Quantitative real-time PCR

Quantitative real-time PCR (qPCR) methods follow those of Feng et al. (2015). Briefly, the qPCR reaction was run on an ABI Viia7 system at the Cornell Genomics Core facility with an annealing temperature of 60.0 °C. Each reaction contained 1ul forward primer, 1ul reverse primer, 5 ul of 2X Power SYBR Green PCR Master Mix (Life Technologies, Carlsbad, CA), 1ul ddH2O, and 2ul of the appropriate cDNA. All cDNA samples were run in triplicate. No template controls were included on each plate. Standard curves for target genes *sod1* and *sod2*, as well as for three reference genes – 18S ribosomal RNA (*18s*), ATP-binding cassette sub-family F member 1 (*abcf1*) and snurportin 1 (*snupn*) were run, in triplicate, on separate plates and used to estimate relative copy numbers of each reaction containing cDNA samples. *Sod1, sod2, 18s, abcf1, and snupn* primers were diluted to a final concentration of 200nM. The appropriate reference genes were chosen for each comparison after optimization experiments (see below). Primer sequences are shown in Table 1.

**Table 1.**
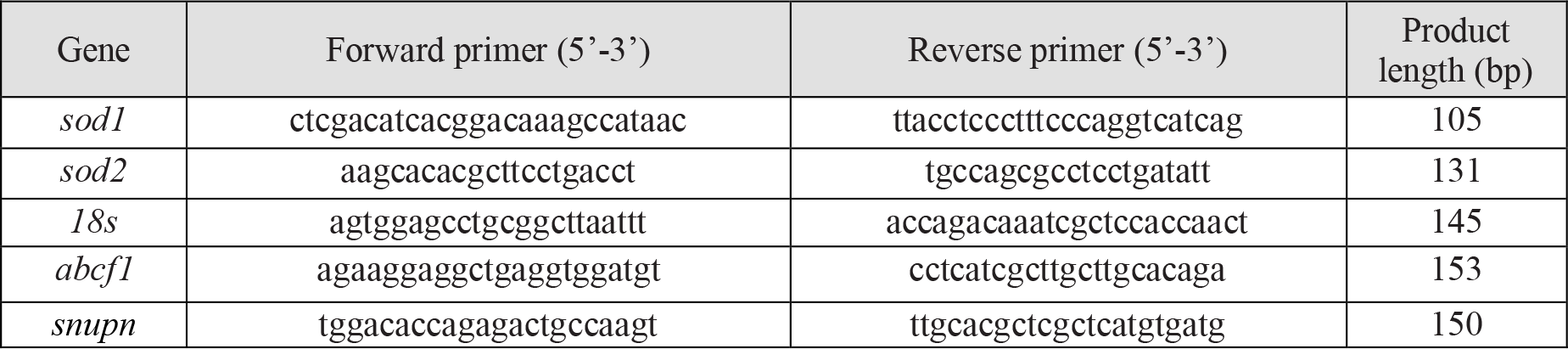
Primer sequences used for qPCR reactions.

### Statistics

All statistical analyses were performed in R v2.1.0 statistical programming language (R Development Core Team (2008). An analysis of variance test (ANOVA) was performed on expression levels of *sod1/18s* and *sod2/18s* in VMN tissue and *sod1/snupn* and *sod2/snupn* in hindbrain tissue to determine if there were statistically significant differences across morphs. In the behavioral state study, *sod1* and *sod2* expression levels were compared between type I males actively humming versus control non-actively humming type I males sacrificed within the same hour. ANOVA was performed on *sod1/abcf1* and *sod2/ancf1* levels in the VMN between the two groups (humming versus control), as well as on *sod1/abcf1* and *sod2/abcf1* levels in the surrounding hindbrain between the two groups. All ANOVA tests were performed on VMN and hindbrain tissue data separately. Assumption of normality was verified with Q-Q plots, visualizing distribution of measurements compared to a normal distribution within each group.

### Selection of Reference Genes

The use of qPCR required that a reference gene be chosen for each analysis that did not vary across the variables of interest: reproductive morph (type I male, type II male, female) in the sex/reproductive study, and actively humming versus non-calling in the behavioral state study. To compare inter-morph variation in the sex/reproductive study, a pilot qPCR study validated 18S as a reference gene that did not vary significantly across morphs in the VMN (ANOVA, F_2,6_ = 0.539, p = 0.609). While *18s* was validated as a reference gene for the VMN, significant *18s* inter-morph variation was found in the SH (ANOVA F_2,26_ = 2.86, p= 0.0754). Therefore, another pilot qPCR study was performed to validate *snupn*, which a transcriptome screen showed has low variance across multiple timescales (summer night, summer morning, winter night), as a suitable reference gene for the hindbrain (ANOVA F_2,6_ = 0.333, p = 0.729) (Feng et al., 2015). Subsequently, qPCR was used to quantify *sod1* and *sod2* expression levels using the standard curve method. Expression levels were examined in the VMN and SH relative to *18s* and *snupn*, respectively. Since we were unable to identify reference genes that showed unvarying patterns of expression across morphs in addition to between VMN and SH tissues, we could only compare *sod1* and *sod2* across morphs for each tissue, but not between tissues (VMN and SH).

To investigate the effect of behavioral state on *sod* levels, a pilot qPCR study first validated *abcf1* as a reference gene that did not vary significantly between humming and non-humming fish in the VMN (ANOVA F_1,8_ = 0.316, p=0.589) as well as in the surrounding hindbrain (ANOVA F_1,10_ = 0.805, p=0.391) (Feng et al., 2015). *Abcf1*, also selected because a transcriptome screen showed it to have low expression variance across multiple timescales (see above), was found to vary between VMN and SH (ANOVA F_1,20_=5.371, p =0.0312) so that we could only make comparisons between behavioral states and not between tissues (VMN and SH).

## RESULTS

### Tissue Comparisons

#### Comparison of reproductive morphs

We observed no significant difference in *sod1/18s* quantity mean across morphs in VMN tissue (ANOVA, F_2,23_ = 0.544, p=0.588) or *sod1/snupn* quantity mean across morphs in the surrounding hindbrain (SH) tissue (ANOVA, F_2,26_ = 1.875, p=0.174) (Fig. 1A, B). Similarly, for *sod2* expression levels, no *sod2/snupn* inter-morph variation was found across morphs in the SH (ANOVA, F_2,26_ = 0.734, p=0.49) (Fig. 1D). However, we did find a significant *sod2/18s* inter-morph variation in the VMN (ANOVA, F_2,23_ = 5.372, p=0.0122). A post-hoc Tukey HSD pairwise comparison was performed, indicating that type I males had a significantly lower mean of *sod2/18s* in the VMN than type II males, but not females (Fig. 1C).

**Figure 1.**
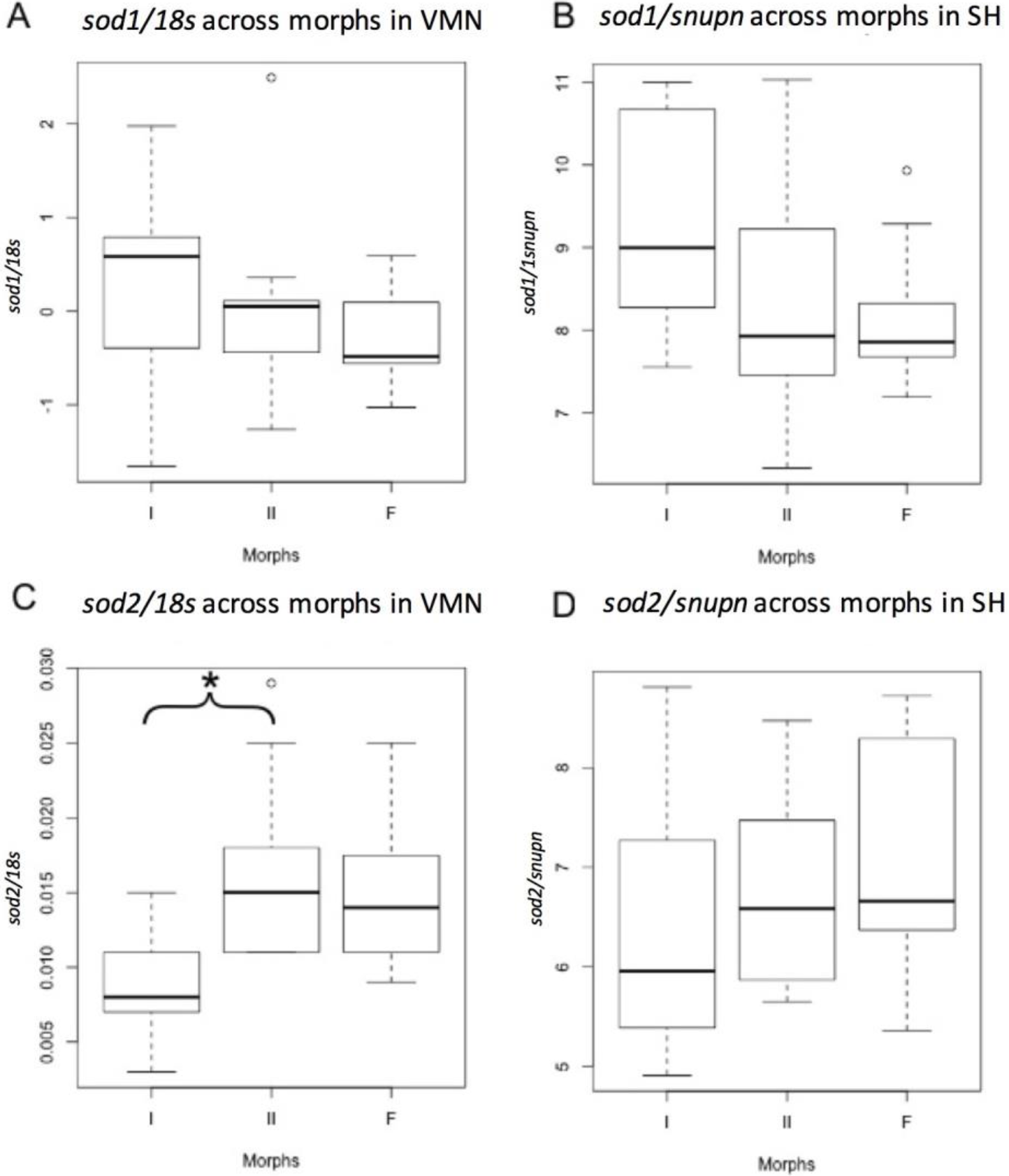
Comparison of *sod1* and *sod2* transcript abundances among the three reproductive morphs of midshipman fish: type I males (I), type II males (II), and females (F). Tissue-specific transcript abundance was compared between morphs using qPCR, and normalized with *18s* for the vocal motor nucleus (VMN) and *snupn* for the surrounding hindbrain (SH). Sample sizes were n=10 for type I males, n=10 for type II males, and n=8 for females. **A.** No significant difference of *sod1* expression across morphs in the VMN. **B.** No significant difference of *sod1* expression across morphs in the SH. **C.** *sod2* was significantly lower in the VMN of type I males as compared to type II males, but not to females. **D.** No significant difference of *sod2* expression across morphs in the SH.

#### Comparison of behavioral state: humming vs non-calling type I males

In the behavioral state study, *sod1/abcf1* levels were found to not vary significantly between humming and non-calling groups in both the VMN (ANOVA F_1,13_ = 1.109, p=0.311) and SH (ANOVA F_1,16_=0.014, p=0.906) (Fig. 2A, B). *sod2/abcf1* levels were also found not to vary significantly between humming and non-calling groups in both the VMN (F_1,12_ = 1.154 p=0.304) and SH (ANOVA F_1,15_ = 0.064, p = 0.803) (Fig. 2C, D).

**Figure 2.**
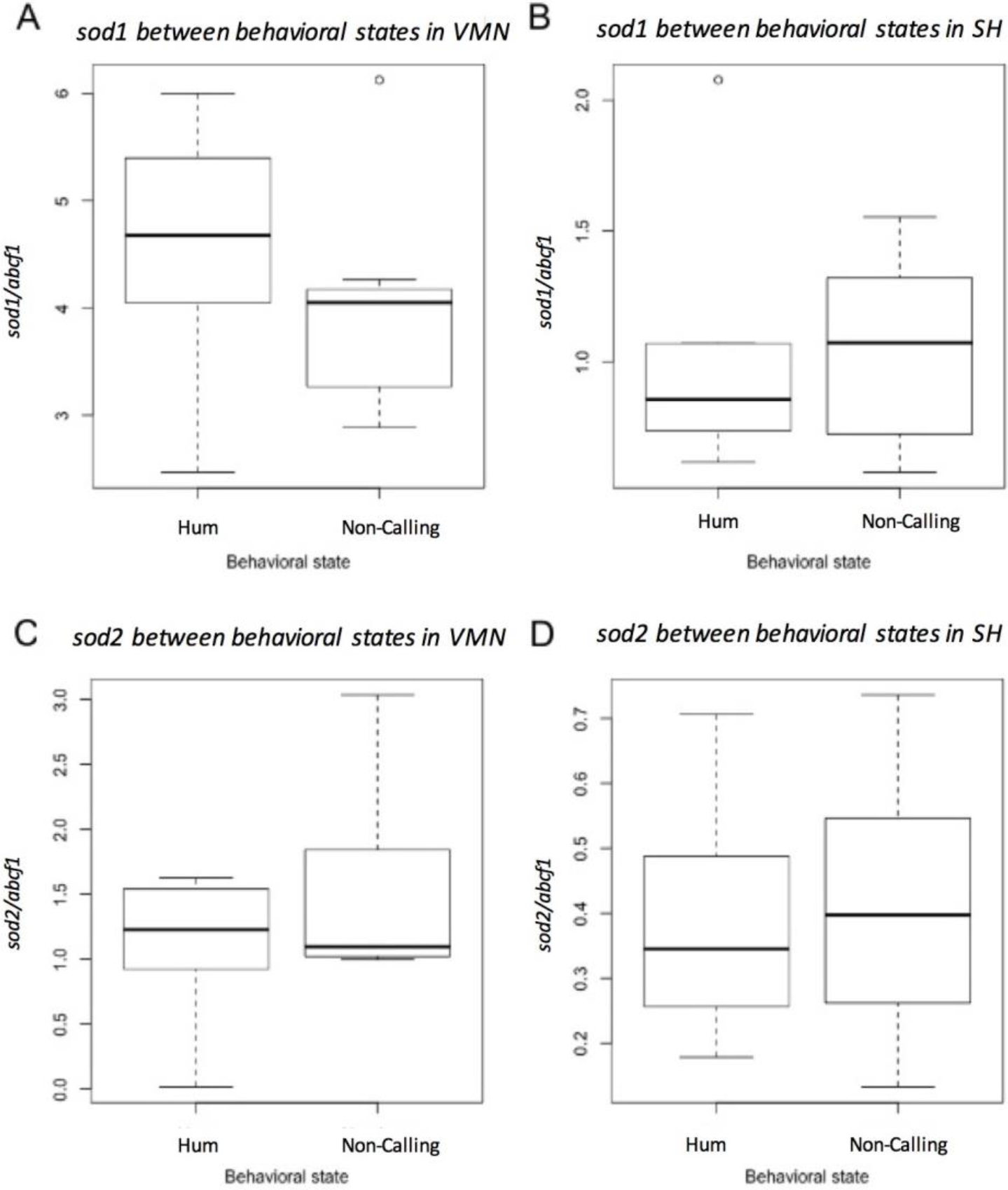
Comparison of *sod1* and *sod2* transcript abundances between actively humming (Hum) and non-calling (Non-calling) type I male midshipman fish. Actively humming fish were allowed to hum for a minimum of 30 min prior to sacrifice followed by immediate extraction of brain tissue. Non-calling fish were anesthetized within the same hour as the actively-humming fish. Tissue-specific transcript abundance was compared between behavioral states using qPCR, and normalized with *abcf1*. **A.** No significant difference of *sod1* expression between humming vs. non-calling groups in the vocal motor nucleus (VMN). **B**. No significant difference of *sod2* expression between groups in the surrounding hindbrain (SH). **C.** No significant difference of *sod2* expression between humming vs. non-calling groups in the VMN. **D.** No significant difference of *sod2* expression between groups in the SH.

## DISCUSSION

In this study, we investigated antioxidant gene upregulation as a potential mechanism for withstanding oxidative stress from potentially high rates of cellular respiration in the midshipman fish vocal hindbrain. More specifically, we tested the hypothesis that to combat oxidative stress, the VMN of type I males that are capable of producing long duration advertisement calls would exhibit higher levels of antioxidant *sod* gene expression compared to the VMN of sneak-spawning type II males and females that are only capable of producing short duration calls.

For animals held in captive conditions not suited for inducing advertisement hum calls in type I males, we found that *sod1* gene expression in the VMN and the SH did not vary significantly across reproductive morphs (type I males, type II males, females), although there was a relatively large variance in expression for both tissues. The large variance led us to investigate the behavioral state dependence of *sod1* expression in type I males, namely asking if *sod1* would be upregulated in type I males actively humming compared to males that were not humming or calling in general. We found, however, no significant difference in *sod1* expression between humming and non-calling type I males in either the VMN or SH. There was also no difference in *sod2* expression across morphs in the SH. In the VMN, however, we observed a significantly lower expression of *sod2* in type I males compared to type II males, although there were no differences between females and either type I or type II males. A companion study showed that *sod2* expression did not vary significantly between humming and non-calling type I males in either the VMN or the SH. Together, the results lead us to conclude that vocal motor neurons in type I males do not show changes in *sod1* and *sod2* gene expression to combat potential oxidative stress imposed by the physiological demands of high-endurance hum vocalizations.

There are multiple potential explanations for the lack of increased transcript expression for the two antioxidant *sod* genes studied here. First, while both genes were among a suite of candidate genes identified to be upregulated in the VMN, it is possible that *sod* expression is constitutively higher in midshipman fish brains, regardless of morph or behavioral state (Feng et al., 2015). Alternatively, other candidate antioxidant genes also showing upregulation in VMN, but not investigated here may show higher expression in type I males as compared to type II males and females, as well as in actively humming compared to non-calling type I males. Additionally, the comparison between the humming and non-calling behavioral states was conducted under the assumption that all VMN neurons were similarly active during humming. If sub-populations drive vocalization during different segments of a hum, or for that matter any vocalization, then differences in expression patterns between them might be hidden by assaying the entire VMN. Last, a well-known caveat when studying mRNA expression is that post-transcriptional mechanisms may alter levels of functional proteins ultimately transcribed, which undermine the assumption that higher RNA levels reflect higher protein levels (e.g., see Gry et al., 2009).

### Steroid-related intra- and intersexual divergence in midshipman fish vocal traits

We found a morph-specific pattern of *sod2* expression that can now be added to the list of intra- (type I versus type II male) and inter-sexual differences in vocal behavior and motor traits (reviewed in Feng and Bass, 2017). To address the surprising result of lower *sod2* expression in type I males compared to type II males in the VMN, we draw insight from other midshipman fish studies. Inspired by neurophysiological studies showing the influence of steroids on vocal motor excitability in midshipman (Remage-Healey and Bass, 2004, 2007), Fergus and Bass (2013) investigated mRNA transcript abundance in VMN for estrogen receptor (ER) genes and aromatase, the enzyme that converts testosterone to estrogen (Bentley, 1998). ERβ2 showed significantly stronger expression in the VMN of females and type II males compared to type I males; ERβ1 showed a similar pattern though the differences were not significant. ERα is significantly higher in type II males compared to type I males and females. Aromatase levels were about two-fold higher in type II males compared to type I males and females, though the differences were not significant. Androgen receptor (AR)α and ARβ expression are also highest in type II males, with overall patterns like those for aromatase and ERβ2, respectively (D. Fergus and A. Bass, unpublished observations). Given the observed dependence of antioxidant enzyme activity in the rat brain on the concentration of progesterone and estrogen (review: Pajovik et al., 2008), we speculate that there may be a functional relationship between steroid signaling and SOD2 levels in midshipman fish. Thus, elevated expression of steroid-related genes in type II males, and in some cases females, compared to type I males may be coupled with enhanced SOD2 levels in the VMN of type II males compared to type I males. In support of this hypothesis, another study showed that the mitochondrial P450-dependent metabolism of steroid hormones activates SOD1 as a protective antioxidant mechanism in liver mitochondria isolated from adult rats, and that both the presence of O_2^−^_ and mitochondrial p450 activity are necessary for the observed SOD1 activation (Inarrea et al., 2011). The potential interaction between steroid hormone pathways and SOD signaling pathways contextualizes the present study’s observed difference in *sod2* expression in the VMN that parallels dimorphic expression patterns in steroid signaling pathways.

### Comparisons with birds and mammals

The endurance challenges that migratory birds face may be discussed as a relevant comparison to those of the midshipman’s courtship humming. Migratory birds have been studied in their ability to increase antioxidants in order to counteract harmful reactive species and oxidative damage. In a recent review on the role of antioxidants in migratory birds, flight was found to have caused oxidative damage in all species examined, but changes in antioxidant enzyme activity during flight varied between species and antioxidant type. For example, glutathione peroxidase was found to be elevated in the red blood cells of European robins (*Erithacus rubecula*) during migratory flight compared to stopover, but unchanging in the Garden warbler (*Sylvia borin*) (Cooper-Mullin and McWilliams, 2016). Zebra finch (*Taeniopygia guttata*) exhibit an unchanging level of SOD during fast flight versus control flight despite an increase in oxidative damage with fast flight, providing an example of a species in which SOD was not the antioxidant to combat oxidative damage (Costantini et al., 2013). This study’s findings of unchanging levels of SOD during high-endurance exercise parallel the present study’s finding of unchanging levels of SOD during high-endurance vocalization. Together, these studies suggest that the pattern of increased levels of antioxidants in tissues facing potential oxidative damage from reactive species produced during high-endurance activity is antioxidant-, tissue- and species-specific.

SOD genes were of particular interest in the prior RNAseq study of midshipman VMN, in part, because of the implications of SOD1 mutations in ALS pathogenesis (Feng et al., 2015). While 90% of ALS cases are known as “sporadic,” with no identified genetic basis attributing to susceptibility, 5-10% of ALS cases are familial. Mutations in SOD1 are observed in 20% of familial ALS cases (Kiernan et al., 2011). ALS pathogenesis is still unclear, but SOD1 has been attributed to some mechanisms of mitochondrial dysfunction leading to motor neuron death (Tafuri et al., 2016). Relative to the our discussion of steroids and sex/morph differences, a potential interaction between hormones and SOD genes is especially intriguing when considering the gender differences observed in ALS. ALS has a higher prevalence in men than women in both familial and sporadic cases (Review: McCombe et al., 2010). Additionally, male mice models with SOD1 mutations show earlier development of ALS than females, suggesting some attribute specific to females possibly hormonal, can delay the onset of the disease (McCombe et al., 2010). A possible correlation between SOD expression and certain hormones may help to explain human gender differences in ALS. It has been proposed that ALS may share some common pathogenic mechanisms related to abnormalities of androgen function, conferring disease susceptibility to ALS (Garofolo et al., 1993). It has also been hypothesized that estrogens serve a protective effect while androgens serve an adverse effect in ALS etiology (Blasco et al., 2012). However, proposed hypotheses on the gender difference in ALS have not been well tested. Studies of SOD expression in fish species like midshipman with alternative reproductive morphs that differ in steroid signaling mechanisms (Brantley et al., 1993; Fergus and Bass, 2013; Remage-Healey and Bass, 2007) can potentially be useful in supplementing research on the role of SOD in not only the pathology but human gender differences in ALS.

### Concluding Comments

The midshipman sonic muscles that vibrate the swim bladder and lead to the production of long duration advertisement hums are known as “one of the most superfast and super-enduring striated muscles found in nature” (Forbes et al., 2006) that have morphological, physiological and molecular adaptations related to this remarkable capacity (Bass and Marchaterre, 1989; Walsh et al., 1995; Forbes et al., 2006; Nelson et al., 2018). It follows that sonic/vocal swim bladder motor neurons would show traits adapted to their ability to drive sonic muscles during such high-stamina contractions. In contrast, other organisms that lack a superfast muscle or the need for such high-endurance motor systems do not necessarily require comparable adaptations to do so. By comparing type I male midshipman that alone are able to generate long duration hums with type II males and females that do not hum, we shed light on the molecular characteristics of motor neurons that allow them to drive superfast muscles, in contrast with other organisms that do not have such muscles (also see Chagnaud et al., 2012).

We chose to investigate *sod1* and *sod*2 transcript expression specifically because a previous RNAseq study showed them to have greater expression in the VMN as compared to the SH in type I males (Feng et al., 2015). To our knowledge, no other vertebrates are known to make individual sounds that last for hours at a time. This ability led us to hypothesize that the midshipman’s unique capability would shed light on the mechanisms that motor neurons have adopted for withstanding high-endurance activity as well as to elucidate information on transcriptional differences in two male reproductive morphs that follow alternative reproductive tactics including humming versus non-humming behavioral states. We have now excluded *sod1*/2 transcript upregulation as a mechanism for protecting vocal motoneurons from oxidative stress during humming, but investigations of other antioxidants are warranted. Narrowing in on the molecular mechanisms responsible for allowing male midshipman to generate long duration vocalizations will help illuminate how different motor neuron populations are adapted to their particular behavioral tasks, e.g., vocalization versus locomotion, as well as the mechanisms behind the comparatively limited capabilities of motor neurons in other organisms.

## Acknowledgments

Thanks to all the members of the Bass Lab for logistical support and to Margaret Marchaterre for collection of the fish used in this study. This research was supported by NSF grant I0S1457108 to Andrew Bass.

## LITERATURE CITED

Adamczyk, B., & Adamczyk-Sowa, M. (2016). New insights into the role of oxidative stress mechanisms in the pathophysiology and treatment of multiple sclerosis. Oxidative Medicine and Cellular Longevity, 2016, 1973834.

Atanasova, S., Wieland, E., Schlumbohm, C., Korecka, M., Shaw, L., von Ahsen, N., & Armstrong, V. (2009). Prenatal dexamethasone exposure in the common marmoset monkey enhances gene expression of antioxidant enzymes in the aorta of adult offspring. Stress, 12(3), 215–224.

Bass AH. (1996). Shaping brain sexuality. American Scientist, 84: 352–63.

Bass, A. H., & Marchaterre, M. A. (1989). Sound-generating (sonic) motor system in a teleost fish (Porichthys notatus): Sexual polymorphism in the ultrastructure of myofibrils. Journal of Comparative Neurology, 286(2), 141–153.

Bass, A. H., & Baker, R. (1990). Sexual dimorphisms in the vocal control system of a teleost fish: morphology of physiologically identified neurons. Developmental Neurobiology, 21(8), 1155–1168.

Bentley, P.J. (1998). Comparative vertebrate endocrinology. Cambridge University Press.

Blasco, H., Guennoc, A. M., Veyrat-Durebex, C., Gordon, P. H., Andres, C. R., Camu, W., & Corcia, P. (2012). Amyotrophic lateral sclerosis: a hormonal condition? Amyotrophic Lateral Sclerosis, 13(6), 585–588.

Brantley, R. K., & Bass, A. H. (1994). Alternative male spawning tactics and acoustic signals in the plainfin midshipman fish Porichthys notatus Girard (Teleostei, Batrachoididae). Ethology, 96(3), 213–232.

Brantley, R. K., Wingfield, J. C., & Bass, A. H. (1993). Sex steroid levels in Porichthys notatus, a fish with alternative reproductive tactics, and a review of the hormonal bases for male dimorphism among teleost fishes. Hormones and Behavior, 27(3), 332–347.

Bruijn, L. I., Miller, T. M., & Cleveland, D. W. (2004). Unraveling the mechanisms involved in motor neuron degeneration in ALS. Annual Review of Neuroscience, 27, 723–749.

Chagnaud, B. P., Zee, M. C., Baker, R., & Bass, A. H. (2012). Innovations in motoneuron synchrony drive rapid temporal modulations in vertebrate acoustic signaling. Journal of Neurophysiology, 107(12), 3528–3542.

Chagnaud, B.P., Baker, R., & Bass, A. H. (2011). Vocalization frequency and duration are coded in separate hindbrain nuclei. Nature Communications, 2, 346.

Cohen, M. J., & Winn, H. E. (1967). Electrophysiological observations on hearing and sound production in the fish, Porichthys notatus. Journal of Experimental Zoology Part A: Ecological Genetics and Physiology, 165(3), 355–369.

Cooper-Mullin, C., & McWilliams, S. R. (2016). The role of the antioxidant system during intense endurance exercise: lessons from migrating birds. Journal of Experimental Biology, 219(23), 3684–3695.

Costantini, D., Monaghan, P., & Metcalfe, N. B. (2013). Loss of integration is associated with reduced resistance to oxidative stress. Journal of Experimental Biology, 216(12), 2213–2220.

Feng N. Y., & Bass A. H. (2017) Neural, hormonal, and genetic mechanisms of alternative reproductive tactics. In Hormones, Brain, and Behavior, Pfaff DW, Joels M (eds), Vol 2, 3rd edn, pp 47–65. Oxford: Academic Press.

Feng, N. Y., Fergus, D. J., & Bass, A. H. (2015). Neural transcriptome reveals molecular mechanisms for temporal control of vocalization across multiple timescales. BMC Genomics, 16(1), 408.

Feng, N. Y., & Bass, A. H. (2016). “Singing” fish rely on circadian rhythm and melatonin for the timing of nocturnal courtship vocalization. Current Biology, 26(19), 2681–2689.

Fergus, D. J., & Bass, A. H. (2013). Localization and divergent profiles of estrogen receptors and aromatase in the vocal and auditory networks of a fish with alternative mating tactics. Journal of Comparative Neurology, 521(12), 2850–2869.

Forbes, J. G., Morris, H. D., & Wang, K. (2006).Multimodal imaging of the sonic organ of Porichthys notatus, the singing midshipman fish. Magnetic Resonance Imaging, 24(3), 321–331.

Garofalo, O., Figlewicz, D. A., Leigh, P. N., Powell, J. F., Meininger, V., Dib, M., Rouleau, G. A. (1993). Androgen receptor gene polymorphisms in amyotrophic lateral sclerosis. Neuromuscular Disorders, 3(3), 195–199.

Gry, M., Rimini, R., Strömberg, S., Asplund, A., Pontén, F., Uhlén, M., & Nilsson, P. (2009). Correlations between RNA and protein expression profiles in 23 human cell lines. BMC Genomics, 10(1), 365.

Ibara, R.M., Penny, L.T., Ebeling, A.W., van Dykhuizen, G., & Cailliet, G. (1983). The mating call of the plainfin midshipman fish, Porichthys notatus. In Predators and Prey in Fishes, pp. 205–212. Doredrecht: Springer.

Inarrea, P., Casanova, A., Alava, M. A., Iturralde, M., & Cadenas, E. (2011). Melatonin and steroid hormones activate intermembrane Cu, Zn-superoxide dismutase by means of mitochondrial cytochrome P450. Free Radical Biology and Medicine, 50(11), 1575–1581.

Kiernan, M. C., Vucic, S., Cheah, B. C., Turner, M. R., Eisen, A., Hardiman, O., … & Zoing, M. C. (2011). Amyotrophic lateral sclerosis. The Lancet, 377(9769), 942–955.

Kumar, A., & Ratan, R. R. (2016). Oxidative stress and Huntington’s disease: The good, the bad, and the ugly. Journal of Huntington’s Disease, 5(3), 217–237.

Lobo, V., Patil, A., Phatak, A., & Chandra, N. (2010). Free radicals, antioxidants and functional foods: Impact on human health. Pharmacognosy Reviews, 4(8), 118–126.

McCombe, P. A., & Henderson, R. D. (2010). Effects of gender in amyotrophic lateral sclerosis. Gender Medicine, 7(6), 557–570.

McIver, E. L., Marchaterre, M. A., Rice, A. N., & Bass, A. H. (2014). Novel underwater soundscape: acoustic repertoire of plainfin midshipman fish. Journal of Experimental Biology, 217(13), 2377–2389.

Nelson FE, Hollingworth S, Marx JO, Baylor SM, Rome LC. (2018). Small Ca+ releases enable hour-long high-frequency contractions in midshipman swimbladder muscle. Journal of General Physiology, 150(1), 127–143.

Pajovic, S. B., & Saicic, Z. S. (2008). Modulation of antioxidant enzyme activities by sexual steroid hormones. Physiological Research, 57(6), 801–811.

Peres, T.V., Schettinger, M. R. C., Chen, P., Carvalho, F., Avila, D. S., Bowman, A. B., & Aschner, M. (2016). Manganese-induced neurotoxicity: a review of its behavioral consequences and neuroprotective strategies. BMC Pharmacology and Toxicology, 17(1), 57.

Pocernich, C.B., La Fontaine, M., & Butterfield, D. A. (2000). In-vivo glutathione elevation protects against hydroxyl free radical-induced protein oxidation in rat brain. Neurochemistry International, 36(3), 185–191.

Power, J.H., & Blumbergs, P. C. (2009). Cellular glutathione peroxidase in human brain: cellular distribution, and its potential role in the degradation of Lewy bodies in Parkinson’s disease and dementia with Lewy bodies. Acta neuropathologica, 117(1), 63–73.

Remage-Healey, L., & Bass, A. H. (2004). Rapid, hierarchical modulation of vocal patterning by steroid hormones. Journal of Neuroscience, 24(26), 5892–5900.

Remage-Healey, L., & Bass, A. H. (2007). Plasticity in brain sexuality is revealed by the rapid actions of steroid hormones. Journal of Neuroscience, 27(5), 1114–1122.

Tafuri, F., Ronchi, D., Magri, F., Comi, G. P., & Corti, S. (2015). SOD1 misplacing and mitochondrial dysfunction in amyotrophic lateral sclerosis pathogenesis. Frontiers in Cellular Neuroscience, 9, 336.

Walsh, P.J., Mommsen, T. P., & Bass, A. H. (1995). Biochemical and molecular aspects of singing in batrachoidid fishes. In Biochemistry and Molecular Biology of Fishes, Vol 4, pp. 279–289. New York: Elsevier.

